# Ligand modulation of allosteric networks in an ancestral steroid receptor

**DOI:** 10.1101/414375

**Authors:** C. Denise Okafor, Eric A. Ortlund

## Abstract

Understanding the evolution of binding specificity, a heavily studied area of research, is key for determining how protein sequence changes alter function. Ligand-activation in the steroid receptor subfamily of transcription factors operates via a common allosteric mechanism which permits extant receptors to respond specifically to their cognate hormones. Here, we combine atomistic simulations with graph theory-based modeling of the inter-residue interactions within protein complexes to gain insight into how allostery drove selectivity in an ancestral receptor. An inactive ligand complex displays weakened allosteric communication, as quantified by suboptimal paths linking two functional surfaces. When function-switching mutations are incorporated, responses in allosteric networks are consistent with ligand activation status. Further analysis reveals residues that modulate features distinguishing active and inactive complexes, identifying a key, conserved residue that is crucial for activation in steroid receptors. We have identified a computational method using dynamic network analysis to probe the allosteric mechanisms driving the evolution of ligand specificity in hormone receptors, determining how residue substitutions altered allosteric networks to permit gain or loss of ligand response. These results may have general utility in elucidating how modern steroid receptors are activated by endogenous and xenobiotic molecules.

**Author summary:** Proteins interact with a host of biological partners to mediate their function. These binding partners are able to alter structural properties of the protein to send signals dictating downstream biological activity. This mode of regulation is described as allostery. Here, we perform a computational investigation of allostery in steroid receptors, a family of proteins that regulate a host of important biological processes in response to binding and activation by a steroidal ligand. We leverage a defined evolutionary system where known historical amino acid substitutions within the receptor drive a switch in ligand preference and receptor activation. We show that activating ligands induce stronger allosteric signaling between the ligand and the functional surface on the receptor. In addition, we incorporate evolutionary mutations that are known to alter ligand preference and show that this effect may be explained by allostery. This work provides insight into how amino acid substitutions over evolution affect allostery in proteins, permitting the loss and gain of function.

## Introduction

Allostery, the biological concept of distant regulation in proteins, has been heavily studied and reviewed (1-3). In its narrowest definition, allostery refers to the phenomenon whereby a molecule binds at a secondary site (i.e. independently of a primary substrate binding site) to regulate protein activity. More broadly, it refers to an observed event at one region of a protein which induces effects at a distant region. Experimentally, these activities were identified structurally, evidenced by conformational changes observed in crystal structures (4). Decades of research have led to an evolution of the characterization of allostery (5, 6). Allostery now encompasses a diversity of established concepts, such as dynamics (7, 8), entropy (9-11), conformational selection (12, 13), and population shift (14, 15), indicative of the increasing inclusivity in our understanding of allostery. An ensemble perspective on allostery (16, 17) accounts for an initial distribution of conformational states within the protein that undergoes a redistribution upon perturbation (e.g. ligand binding), reflective of the transmission of an allosteric signal.

An allosteric network describes the concerted sets of interactions that are most implicated in signal propagation between coupled regions of a protein (18). Often considered to be pathways formed by residues mediating communication between the two sites, these networks have been identified in a variety of ways. Experimental techniques used to identify allosteric networks include nuclear magnetic resonance (NMR) (19), hydrogen-deuterium exchange mass spectrometry (HDX-MS) (20, 21), x-ray crystallography (22, 23), and anisotropic thermal diffusion (24). Additionally, molecular dynamics (MD) simulations in combination with a range of specialized analytical methods have been used to elucidate plausible protein allosteric networks (25). Theoretical methods have also identified allosteric networks. These include graph or network theory (26), often used in combination with MD (27), and statistical coupling analysis (SCA), utilizing energetic evaluation of evolutionarily conserved residues within protein families (28).

The activity of the Nuclear Receptor (NR) subfamily of transcription factors can be characterized by long-range coupled interactions (29). NRs are multidomain receptors that regulate gene expression in response to ligand binding (30). Common NR domains are the N-terminal domain (NTD), which contains an activation region; the DNA binding domain (DBD), which binds DNA response elements; and the ligand binding-domain (LBD), which binds small hydrophobic molecules with high specificity. These lipophilic ligands dock in a hydrophobic pocket within the LBD, initiating effects that propagate to multiple intra-domain regions within the LBD, including a dimerization interface and coregulator binding surface (also known as the activation function surface 2, AF-2) (31, 32). Coregulators interacting at the AF-2 surface fall into two main categories: coactivators which promote transcriptional activity and corepressors which repress transcription (33). In addition, inter-domain allosteric effects are observed. Ligand binding can regulate the interactions between NR domains and a host of molecular binding partners (34, 35) and the LBD is subject to modest regulation by the DBD in a DNA sequence specific manner (36, 37).

The modular nature of NRs permits ease of structural and biochemical characterization of isolated domains. Experimental analysis of isolated NR LBDs has allowed for the elucidation of ligand specificity, and in many cases the generation of high-affinity small molecule modulators. Steroid Receptors (SRs) are a subclass of NRs (subfamily III) that modulate transcription of a host of genes specifically in response to binding of steroidal hormones. This subfamily consists of estrogen receptors (ERα and ERβ), glucocorticoid receptor (GR), mineralocorticoid receptor (MR), progesterone receptor (PR) and the androgen receptor (AR).

The SR family is comprised of two major clades (Figure 1) (38). The first clade contains the ERs which are activated by estrogenic hormones, i.e. steroids with an aromatized A-ring and a hydroxyl at the carbon 3 (C3) position (Figure 1). The second clade contains the non-aromatized steroid receptors (naSRs): GR, AR, MR, and PR, all of which respond to hormones with a non-aromatized A-ring, possessing either a keto or a hydroxyl group at the C3 position (39). Ancestral sequence resurrection (ASR) is a powerful phylogenetic technique that has been used to infer ancestral protein sequences (38, 40), permitting their resurrection and subsequent characterization. This approach has been used to define the evolution, expansion and subsequent functional specification of the SR family (38, 41). The first SR is termed ancestral steroid receptor 1 (AncSR1), emerged ∼550 million years ago (mya) (41, 42). AncSR1 was estrogen-sensitive, but unresponsive to all steroids with a non-aromatized A-ring, and thus is classified as estrogen receptor-like (39).

**Figure 1:**
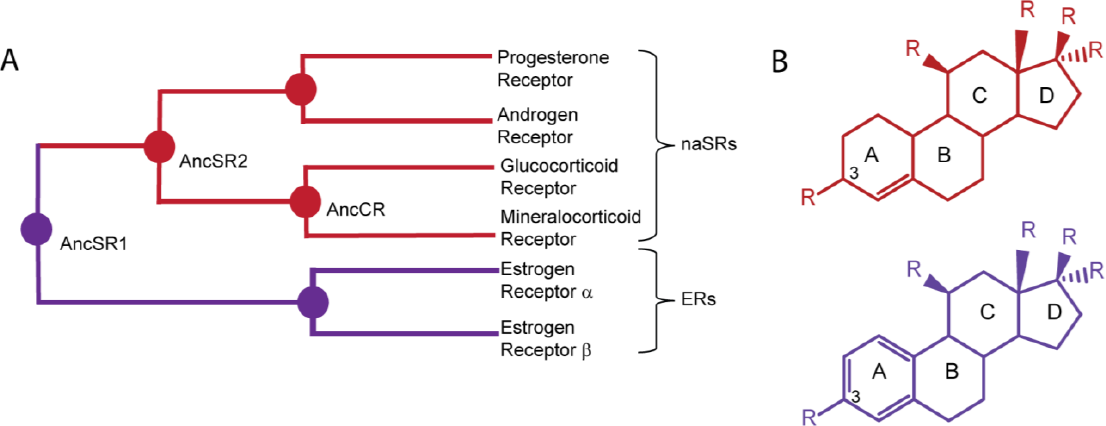
Steroid receptors evolved from an estrogen-responsive ancestor. A. Simplified phylogeny of the steroid receptor family. Two clades comprise the SR family, non-aromatized steroid receptors (naSRs) and estrogen receptors (ERs). B. Steroid hormone scaffold with rings and C3 position labeled. Non-aromatized A-ring scaffold is shown in red, while the aromatized A-ring scaffold is shown in purple.

The ancestral steroid receptor 2 (AncSR2) is the ancestor of all naSRs, loses all response to aromatized steroids (e.g. estrogens) and was shown to be responsive to a broad range of non-aromatized steroids, including androgens, progestogens, and corticoids (42). Subsequent ASR studies have investigated evolutionary causes of both promiscuity and specificity across SRs, as well as the specific sequence changes that have permitted modern SRs to retain responsiveness to their cognate hormones while losing activation from other hormones (43-46). For example, two residue substitutions in AncCR (Figure 1), S106P and L111Q, were shown to recapitulate the GR functional shift, reducing activation by aldosterone and increasing selectivity for cortisol (47).

Allosteric networks in extant SRs have been described between the LBP and AF-2 surface (31, 48-50). These studies have shed light on ligand-coregulator specific effects of both endogenous steroid hormones and synthetic SR ligands. Allosteric networks have not been investigated in ancestral SRs, so it is not clear how these networks have evolved in concert with ligand specificity, to permit the gain and loss of hormone recognition. A semi-promiscuous receptor such as AncSR2 presents an ideal model in which to investigate allosteric networks induced by different activating ligands, as well as to examine how non-activating ligands alter the network in way that is not productive for coactivator recruitment and transcriptional activation.

Here, we investigate allosteric networks in AncSR2 in complex with steroid hormones from each receptor class: androgens (dihydrotestosterone), progestogens (progesterone), mineralocorticoids (aldosterone), glucocorticoids (cortisol), and estrogens (estradiol) (Figure 2). With a goal of identifying the allosteric effects that accompany SR activation, we focus on detecting features that distinguish the non-activating complex (i.e. AncSR2-estradiol) from the activating complexes.

**Figure 2:**
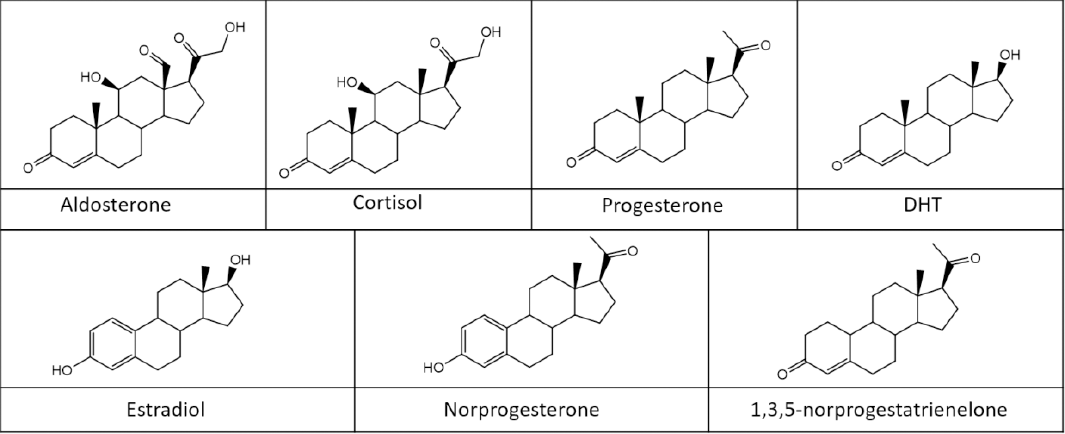
Steroid hormones in this study.

Next, we examine the effects of amino acid mutations on AncSR2 allosteric networks. Two historical AncSR1 amino acid substitutions were identified which upon insertion in AncSR2, reverse the evolutionary shift in hormone specificity. Two steroids, progestagen norprogesterone (NOR) and the estrogen 1,3,5-norprogestatrienolone (NPT) were selected because they are identical except that NOR is a nonaromatized 3-ketosteroid while NPT is aromatized with a 3-hydroxyl (Figure 2). AncSR2 was shown to be responsive to the progestagen norprogesterone (NOR) but weakly responsive to the estrogen 1,3,5-norprogestatrienelone (NPT) (Figure 2), whereas AncSR1 shows activation by NPT but not NOR. Reverse amino acid substitutions Q41E and M75L in AncSR2 dramatically increased sensitivity to NPT and reduced activation by NOR.

To describe allosteric networks, we utilize network theory in combination with MD simulations (51). Networks represent the protein as a collection of nodes and edges. Nodes are defined at each Cα atom, and edges are drawn to connect the nodes if the residues are within a pre-defined cutoff for the simulation. Edges are weighted by selected atomic properties, and paths can be drawn between distant nodes in the network by connecting them with a chain of edges. Therefore, network analysis provides insight into the mechanisms by which non-covalent contacts lead to long-range communication in proteins. In addition to allostery and molecular signaling, network analysis has been used to investigate many features of protein structure and function, such as protein folding, conformational analyses and protein-ligand binding (52, 53).

Our results show allosteric networks in AncSR2 are differentially modified by distinct ligands, depending on their activating properties. Key amino acid substitutions modulate allosteric networks in a ligand-dependent fashion as well, allowing for correlations to be drawn between receptor activity and allosteric communication quantified by path lengths. These results imply that the dynamics of AncSR2 complexes are intrinsically different in the presence of different bound ligands. Even without large conformational shifts, allosteric networks clearly reflect these distinctions, permitting investigation into how steroid receptors evolved strong ligand specificity for their cognate steroid hormones, despite high structural similarity.

## Methods

### Structure preparation

AncSR2 LBD-ligand complexes were prepared for molecular dynamics simulations, constructed from PDB 4LTW (54) which shows AncSR2 bound to progesterone in the ligand binding pocket (Figure 3). All waters and surface-bound molecules derived from the crystallization buffer were deleted from AF-2, so that complexes consisted of AncSR2 and steroidal ligands. All modifications were introduced *in silico* to the modified PDB 4LTW.

**Figure 3:**
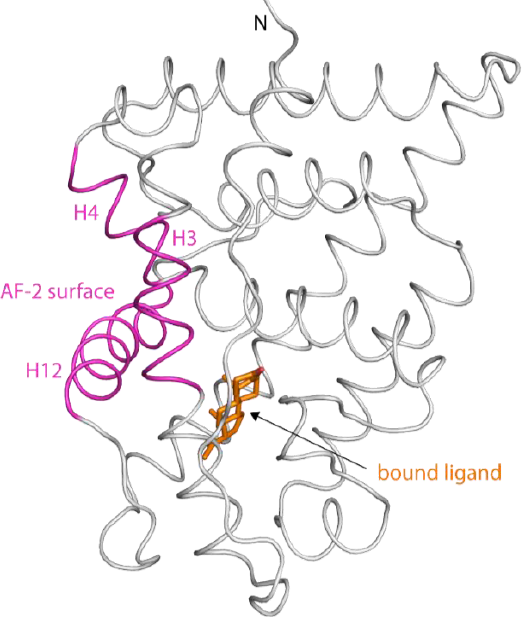
Cartoon representation of the AncSR2 structure (PDB 4LTW). Progesterone ligand (orange) is in the pocket (LBP). The activation function 2 (AF-2) surface, colored magenta, is comprised of helices 12, 3 and 4 (H12, H3, H4) and is the site for binding to transcriptional co-regulatory proteins required for gene activation.

Modifications were made in two sets:

Set A: Five AncSR2 complexes were constructed from 4LTW by modifying the progesterone core to obtain the following hormonal complexes: i) AncSR2-aldosterone; ii) AncSR2-cortisol; iii) AncSR2-progesterone; iv) AncSR2-dihydrotestosterone (DHT) and v) AncSR2-estradiol.

Set B: Ten AncSR2 complexes were constructed from 4LTW by introducing mutations to wild-type AncSR2, as well as modifications to the progesterone scaffold to obtain either NOR or NPT. The following complexes were constructed: i) AncSR2-NOR, ii) AncSR2-NPT, iii) Q41E-AncSR2-NOR, iv) Q41E-AncSR2-NPT, v) M75L-AncSR2-NOR, vi) M75L-AncSR2-NPT, vii) Q41E/M75L-AncSR2-NOR, viii), Q41E/M75L-AncSR2-NPT, ix) Q41E/M75L/R82A-AncSR2-NOR, x) Q41E/M75L/R82A-AncSR2-NPT.

### MD simulations

The complexes were solvated in an octahedral box of TIP3P water with a 10-Å buffer around the protein. Na^+^ and Cl^-^ ions were added to neutralize the protein as well as achieve physiological concentrations. All systems were set up using xleap in AmberTools 17, contained in Amber 2017 (55). All minimizations and simulations were performed with Amber 16, also contained in Amber 2017.

The minimization protocol was as follows: 5000 steps of steepest descent followed by 5000 steps of conjugate gradient minimization with i) 500-kcal/mol.A^2^ restraints on protein and ligand atoms; ii) 100-kcal/mol.A^2^ restraints on protein and ligand atoms, iii) 100 kcal/mol.A^2^ restraints on the ligand atoms, iv) no restraints.

All complexes were heated from 0 to 300 K using a 100-ps run with constant volume periodic boundaries and 5-kcal/mol.A^2^ restraints on all protein and ligand atoms. MD equilibration was performed in the following stages: i) 10 ns with 10-kcal/mol.A^2^ restraints on protein and ligand atoms using the NPT ensemble, ii) 10 ns with 1-kcal/mol.A^2^ restraints on protein and ligand atoms, and iii) 10 ns with 1-kcal/mol.A^2^ restraints on ligand atoms. Restraints were then removed and 1 microsecond production simulations were performed for each system in the NPT ensemble.

A 2-fs time step was used, with all bonds between heavy atoms and hydrogens fixed with the SHAKE algorithm (56). A cutoff distance of 10 Å was used to evaluate long-range electrostatics with particle mesh Ewald and van der Waals forces. Structural averaging and analysis were performed with the CPPTRAJ module of AmberTools17 (57). The ‘strip’ and ‘trajout’ commands of CPPTRAJ were used to remove solvent atoms and obtain fifty-thousand evenly spaced frames from each simulation for analysis. Hydrophobic interactions were defined between ligand and protein C atoms if the atomic distance was < 6 Å for 75% of the simulation. The ‘distance’ command of CPPTRAJ was used to obtain these distances.

### Network Analysis

The NetworkView plugin in VMD (58) and the Carma program (59) were used to produce dynamic networks for each system. Networks are constructed by defining all protein C-α atoms as nodes, using Cartesian covariance to measure communication within the network. An edge is drawn between every pair of non-sequential nodes that reside within 4.5 Å for 75% of the simulation. Edges are weighted by their correlated motion in the MD trajectory, such that edge weights are inversely proportional to the covariance between the nodes. To generate networks, all solvent atoms were stripped, leaving ligand and protein atoms, and the protocol described in (51) was followed. Carma also allows the generation of covariance matrices over C-α atoms.

### Suboptimal Paths

Strength of communication between the ligand and activation function helix was assessed by examination of suboptimal paths between these sites. While communication between two regions of a receptor can occur between thousands of possible paths, suboptimal paths analysis predicts routes and strengths of allosteric communication. A communication path can be drawn as a chain of edges connecting the two nodes. Due to the inverse correlation between correlation and edge weights, the sum of edges along a path between two distant nodes becomes lower as the strength of communication (i.e. correlation) increases. The optimal path is defined as the path for which the sum of edges is the lowest. However, a set of the shortest suboptimal paths, along with the optimal path, are thought to convey the greatest amount of communication between two distant nodes. The Floyd-Warshall algorithm (60) was used to identify suboptimal paths. In this study, all suboptimal paths are calculated between the bound ligand located in the ligand binding pocket (LBP) and helix 12 (H12), which is a component of AF-2, the site of coregulator binding on the LBD (Figure 3).

## Results

### AncSR2-hormone complexes

#### Suboptimal path analysis between ligand and AF-2

AncSR2 is activated by progesterone, DHT, aldosterone and cortisol, but not by estradiol. To describe and quantify allosteric signaling between the hormones and H12/AF-2, we identified the shortest 2000 suboptimal paths connecting the two sites in all five AncSR2-hormone complexes. We observe that the distribution of path lengths is similar between all activating ligands. However, the non-activating complex (SR2-estradiol) displays a distribution with higher path lengths. Therefore path length appears to be correlated with the strength of signaling, and longer path lengths in AncSR2-estradiol are indicative of weakened allosteric communication in AncSR2-estradiol, compared to activating complexes (Figure 4). This weakening could be between the two sites (ligand and H12/AF-2) or possibly resulting from weakened networks throughout the inactive complex.

**Figure 4:**
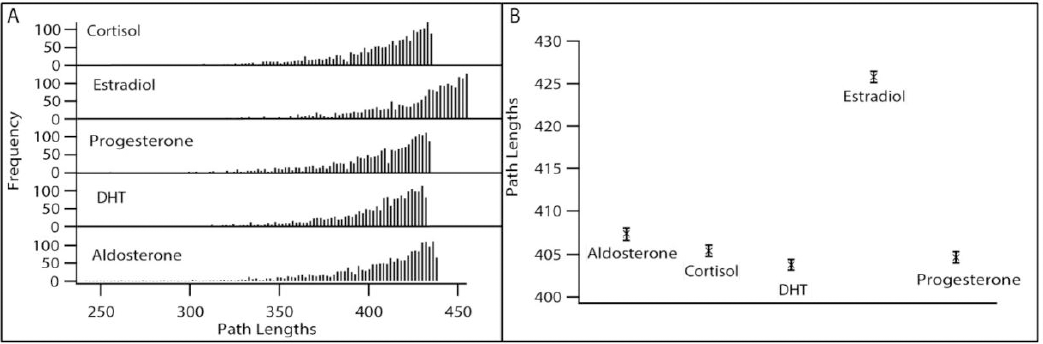
Allosteric signaling between ligand and H12 is weakened in AncSR2-estradiol. A. Histograms show the 2000 shortest suboptimal paths to quantify strength of allosteric signaling in AncSR2-ligand complexes. AncSR2-estradiol shows the weakest (i.e. longest) distribution of paths. B. Average of the 2000 shortest suboptimal paths is plotted for each ligand (Error bars, SEM).

#### Edges implicated in ligand-H12 communication

Communication pathways are comprised of edges that display the strongest correlation throughout the simulation. For each of the AncSR2-hormone complexes, we sought to identify the edges that are implicated in the top 2000 suboptimal paths, with the hypothesis that these edges would reveal the pairwise-residue interactions that are important for allosteric communication. Paths are drawn between the bound hormone (Figure 5A, orange) and a H12 residue (Figure 5A, magenta). We have summarized this set of top edges and the frequency with which they appear in the AncSR2-hormone complexes in Table S1. The table contains all edges which appear in at least 15% (i.e. in at least 300 of the top 2000) of the suboptimal paths for any of the AncSR2-hormone complexes.

**Figure 5:**
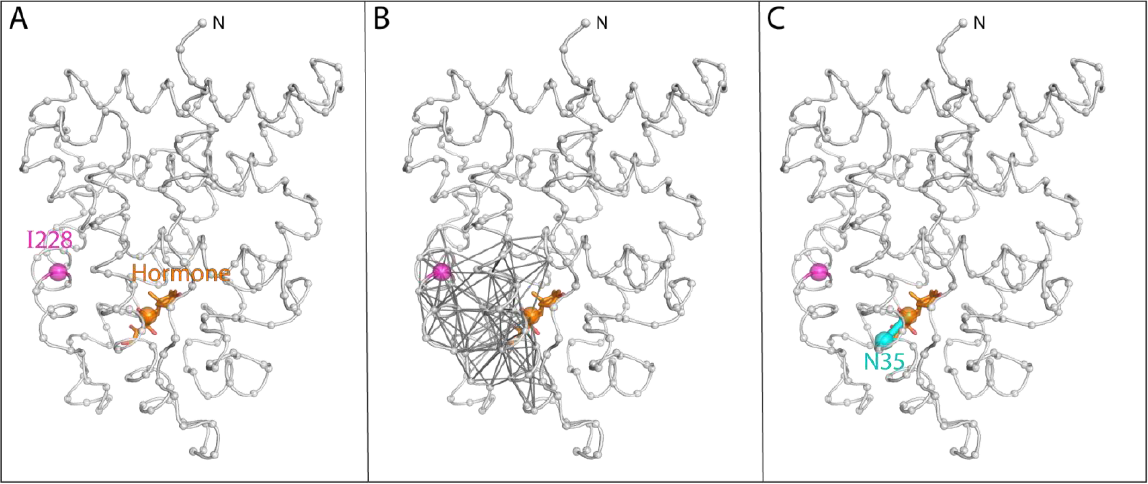
Residues in AncSR2 allosteric signaling. A. Two end-points were selected for quantification of suboptimal paths: C11 atom on hormone and Cα atom of I228 on H12. Spheres indicate positions of selected atoms. B. Suboptimal paths (i.e. chains of edges) connecting bound hormone with I228. C. The edge between the hormone and N35 on H3 (Ligand-N35) occurs with the highest frequency in all AncSR2 complexes studied here.

We observe that the hormones display slight variations in suboptimal path preferences, indicative of ligand-specific patterns of correlation between edges in the complexes (e.g. Figure 5B shows paths for AncSR2-cortisol complex). However, we observe that one edge (ASN35-hormone) is dominant, with high frequency (> 30%) in all five complexes (Figure 5C, cyan). This observation suggests that this edge is important in ligand-H12 communication in all of the complexes studied here.

#### Interactions of hormone in the pocket

The steroid receptor ligand binding pocket is well characterized, with a known set of residues stabilizing the bound hormone via both polar and non-polar interactions (Figure S1). A water molecule in the binding pocket participates in a ligand-protein hydrogen bonding network and is a ubiquitous feature in AncSR LBD-ligand crystal structures. In our simulations, we observe that water molecules penetrate the cavity and integrate into the hydrogen bond networks near the A-ring. In all simulations, the hormone makes expected polar interactions with Arg82, Gln41, Cys207, Thr210 and Asn35 (Figure S1).

To further investigate the observed reduced allosteric strength in AncSR2-estradiol, we compared the binding pocket interactions of estradiol and DHT, two ligands that are identical except for the A-ring (Figure 2). We measured C-C distances (C_ligand_-C_protein_) over the trajectory to identify hydrophobic interactions that persist in the simulation. To quickly highlight the differences between the two ligands, we removed all interactions that were common to the two and show the net differences (Figure 6). This analysis reveals that the two hormones are tilted differently in the pocket. DHT feels a stronger pull on C and D rings towards AF-2, by residues on H3, H4 and H12 (Figure 6A, C, cyan). Conversely, estradiol is pulled on the A-ring end towards H5 and beta strands S1/S2 in the opposite direction (Figure 6C, green), indicating that the ligands sit differently in the pocket. F94, a conserved residue in the SR family, forms a pi-pi interaction with the A-ring that persists over the simulation.

**Figure 6.**
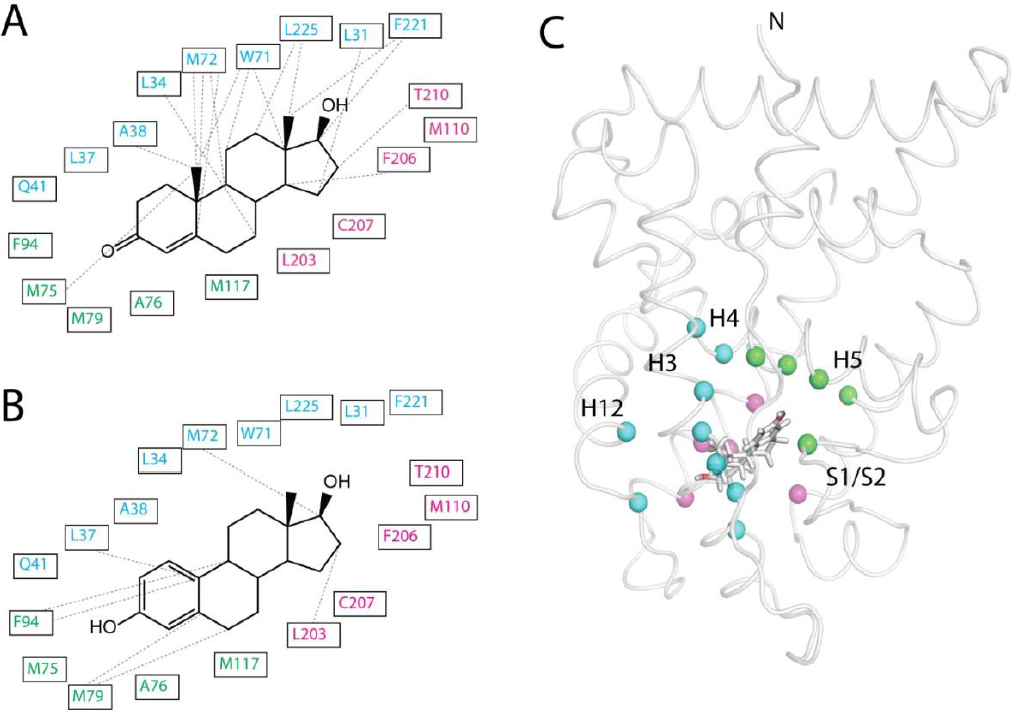
Binding pocket interactions differ between ligands. A-B. Hydrophobic interactions within the pocket differ between DHT (A) and estradiol (B). Dashed lines show residues for which additional hydrophobic interactions (measured as C_ligand_-C_protein_ < 6 Å) were identified, indicative of a tilt in the ligand towards those residues. C. Residue positions indicated in A-B highlighted on AncSR2. The protein is shown as a white loop with Cα atoms represented as spheres. Binding pocket residues are clustered into three groups by location: H5 and S1/S2 beta strands (green), H3/H4/H12 (cyan) and H10/H7 (magenta).

#### AncSR2 with function-switching mutations

Combined phylogenetic and structural studies identified two large-effect substitutions (out of 171 amino acid differences) that explain the difference in hormone specificity between AncSR1 and AncSR2. These two A-ring contacting residues, positions 41 and 75, were seen to dictate hormone preference. The historical reversal of these two residues in AncSR2 (GLN41GLU and MET75LEU) revealed them to be large-effect mutations that drastically shifted AncSR2 hormone preference to the ancestral AncSR1 phenotype.

The MET75LEU substitution significantly increased AncSR2’s affinity for estrogens, measured by luciferase reporter assays, while the GLN41GLU substitution yielded both increased affinity for the estrogenic NPT with reduced sensitivity to the non-aromatized steroid, NOR (61). The double-mutant AncSR2 variant displayed dramatic increases in estrogen sensitivity, accompanied by reduced sensitivity to 3-ketosteroids amounting to a 70,000-fold increased preference for estrogens over a non-aromatized analog. Additionally, introducing the derived states in AncSR1 (i.e. GLU41GLN/LEU75MET) reduced estrogen sensitivity and increased preference for non-aromatized steroids (61). Following this experimental strategy, we reversed these historical amino acid substitutions individually and in combination in AncSR2 (i.e M75L-AncSR2, Q41E-AncSR2, Q41E/M75L-AncSR2) to examine how the estrogenic NPT and non-aromatized NOR organize different allosteric networks in the receptor. The B, C and D rings of the NPT and NOR are identical (Figure 2), allowing us to probe the effect of the aromatized vs the non-aromatized-A ring.

#### Suboptimal path analysis between ligand and H12

We sought to identify how the ancestral substitutions, separately and combined, would alter allosteric signaling in AncSR2-hormone complexes. We begin by quantifying allosteric signaling between the hormone and H12, using the shortest 2000 suboptimal paths connecting the two sites. In AncSR2 complexes with the non-aromatized NOR, we observe that reversing the historical substitutions weakens ligand-H12 communication by increasing path lengths. A strong effect is not seen with the individual mutations, but the response is pronounced in the combined mutant Q41E/M75L-AncSR2 (Figure 7).

**Figure 7:**
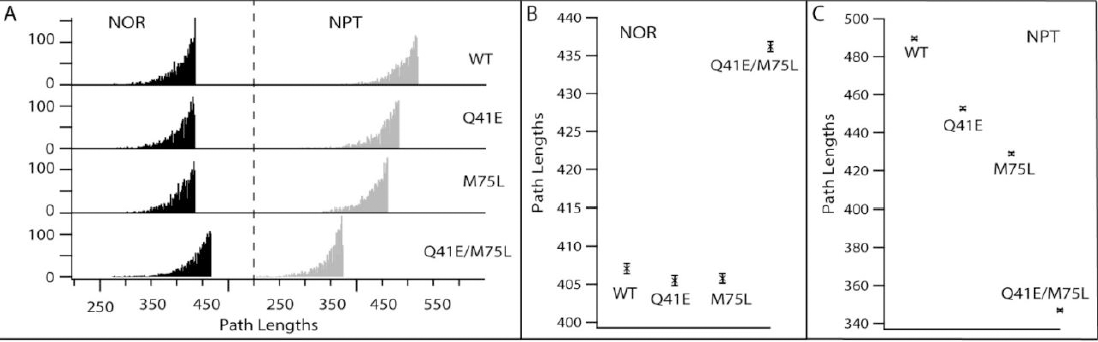
Evolutionary substitutions weaken allosteric signaling in AncSR2-NOR but strengthen signaling in AncSR2-NPT. A. Histograms show the 2000 shortest suboptimal paths to quantify strength of allosteric signaling in AncSR2-NOR (black) and AncSR2-NPT (grey) complexes. WT represents AncSR2 with no residue substitutions. Evolutionary substitutions Q41E and M75L weaken allosteric signaling in NOR complexes, increasing path lengths (black). Conversely, substitutions strengthen signaling in NPT complexes, shifting distributions to shorter path lengths (grey). B-C. Average of the 2000 shortest suboptimal paths is plotted for AncSR2-NOR (B) and AncSR2-NPT (C) complexes showing changes upon incorporation of residue substitutions (Error bars, SEM).

In AncSR2 complexes with the estrogen NPT, ancestral substitutions result in stronger ligand-H12 communication. Individual substitutions (M75L-AncSR2 and Q41E-AncSR2) both lead to shorter suboptimal path lengths, but the combined substitutions further enhance the response, indicative of stronger allosteric networks. These results indicate that the historical substitutions modulate allosteric communication in AncSR2 in a hormone-specific manner, consistent with experimental characterizations of hormone specificity in luciferase reporter assays. Additionally, we performed a deeper analysis of the edges in the top 2000 suboptimal paths for each NOR and NPT complex summarizing the top edges and their frequencies in Table S2, showing the edges that appear in at least 15% of the paths for any of the hormone complexes. This analysis shows once again the dominance of the ASN35-hormone edge (Fig 5C), which is present in high frequency in all 8 complexes. This result confirms the importance of this interaction in ligand-AFH communication in AncSR2, even in the presence of historical substitutions.

#### Effect of historical function-shifting substitutions on networks

Our results suggest that allosteric signaling is altered upon the introduction of evolutionary substitutions in a hormone-specific manner. To identify the subtle changes that occur in the AncSR2 complexes investigated here, we examine covariance plots of mutated AncSR2-complexes and compare them with wildtype AncSR2 (Figure 8). This analysis allows us to visualize changes in correlation that might explain altered allosteric communication in the complexes. *NOR complexes*: Q41E and M75L mutations both induce patterns of anti-correlation in the receptor at similar sites. The double-mutant displays similar patterns as well. Most notably, anti-correlation increases between H3 and multiple regions including H12, H7, H8 and H4/H5. Anti-correlation between H4/5 and H10 increases also (Figure 8, top row). This analysis indicates that weakened ligand-H12 communication could result from changes in correlation throughout the protein, as opposed to one localized region. *NPT complexes*: in contrast to NOR complexes, the introduction of mutations results in the loss of small regions of anti-correlation throughout the complex. M75L and Q41E reveal similar effects, but the double mutant introduces strong anti-correlation between the bottom of H3 and H12, H4/5 (Figure 8, bottom row). These less pronounced effects on correlation observed in AncSR2-NPT complexes emphasize that large allosteric effects can result from subtle changes in conformational dynamics.

**Figure 8:**
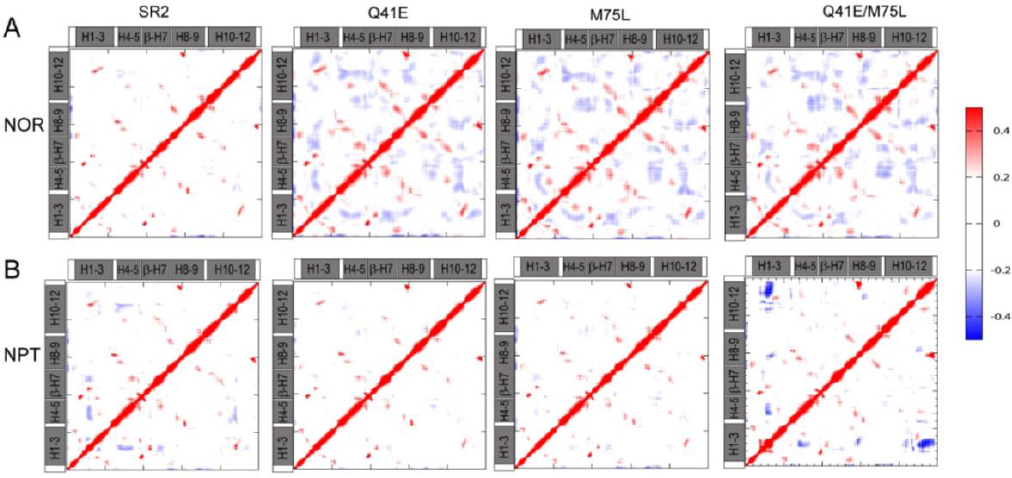
Evolutionary substitutions increase anti-correlation in AncSR2-NOR complexes. Cross-correlation matrices show correlated (red) and anti-correlated (blue) motion over AncSR2 complex simulations. Helical regions are labeled on plots. Four AncSR2 variants (AncSR2, Q41E-AncSR2, M75L-AncSR2 and Q41E/M75L-AncSR2) in complex with A. NOR and B. NPT ligands are shown. Addition of substitutions increases anti-correlation in NOR complexes (top), indicating that global effects in the protein contribute to weakened signaling observed between ligand and H12 Addition of individual substitutions have less pronounced effects on NPT complexes.

#### A mutation that obliterates ligand-specificity erases differences in networks

Arginine 82, a conserved residue in the binding pocket of all SRs, contributes to the hydrogen bonding network with the ligand, interacting with both the C3 group on the A-ring and the residue at position 41 (Figure S1). It was hypothesized that Arg82 plays a special role in mediating the effects of the two historical substitutions, by providing an excess of hydrogen bond donors near the A-ring. To test the hypothesis, Harms et al mutated R82 to A82, eliminating the potential hydrogen bond donors provided by the arginine side chain (61). This substitution removed the dramatic shift in ligand specificity that was observed with the two mutations. The ∼70,000-fold change in hormone preference dropped to a mere 4-fold difference. Thus, reduction in the availability of hydrogen bond donors at the A-ring of the ligand served to diminish the effect of altering function-shifting residues.

We introduced the R82A mutation into MD simulations to determine whether allosteric networks would also reflect the loss of A-ring adjacent hydrogen bond donors. Consistent with experiments, the difference in suboptimal path lengths is greatly reduced (Figure 9). In the double-mutant M75L/Q41E-AncSR2 variant, the NPT complex displays shorter communication paths than the NOR complex, consistent with the idea that reverse evolutionary substitutions increase preference for the estrogen over the non-aromatized steroid. However, when R82A is incorporated into the double mutant background, the loss of hydrogen bond donors appears to mitigate the effects of the historical substitutions, resulting in similar length distributions in the top 2000 paths of both hormone complexes. This result suggests that Arg82 differentially modulates allosteric networks between NOR and NPT complexes, driving ligand-selective effects of the historical substitutions.

**Figure 9:**
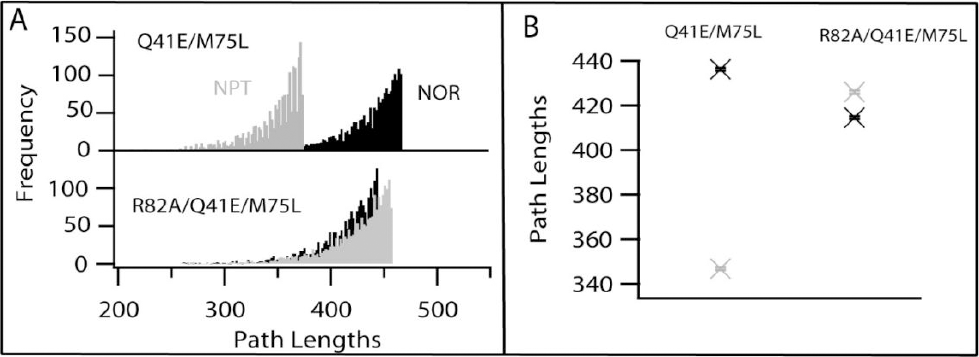
Historical substitutions require R82 to mediate ligand-selectivity. Histograms show the 2000 shortest suboptimal paths to quantify strength of allosteric signaling in Q41E/M75L-AncSR2-NOR (black) and Q41E/M75L-AncSR2-NPT complexes (grey) before (A) and after (B) the introduction of R82A. Addition of R82A to the double AncSR2 mutant (Q41E/M75L-AncSR2) results in markedly different responses to NPT and NOR ligands. The incorporation of R82A reduces the difference in responses between the ligands. (Error bars, SEM).

## Discussion

### Allosteric networks are modulated by ligands and mutations

Steroid receptors operate via an allosteric mechanism in which ligand binding alters the conformational state of the AF-2 region to drive the recruitment of proteins involved in gene expression. Specifically, agonists stabilize AF-2, promoting coactivator binding, while antagonists induce displacement of H12, preventing coactivator binding and/or promoting corepressor binding (62-64). Therefore, in examining allosteric communication between the LBP and H12 in AncSR2, our prediction was that the networks could distinguish between activating and repressing ligands. This difference between active and inactive complexes emerged in the form of longer edges in active complexes, resulting in overall weaker suboptimal paths. The inherent dynamics of the complex in the simulation appear to reflect the ‘inactive’ status of the ligand, resulting in a separate clustering of active complexes. We observe this is also true when AncSR2 is complexed with synthetic estrogen NPT. Consistent with a weak estrogen response in AncSR2 (61), the longest suboptimal paths are observed in AncSR2-NPT.

Furthermore, we observe that the networks generated in MD simulations respond to the incorporation of *in silico* mutations in a manner consistent with experimentally-determined receptor-ligand preferences. Two mutations restore an ancient, AncSR1-like phenotype to AncSR2 by increasing response to estrogen and reducing activation by non-aromatized steroids. When M41L and Q75E are introduced to AncSR2, suboptimal paths were shortened in the estrogen complex but lengthened (i.e. weakened) in the NOR (non-aromatized steroid) complex. Therefore, the effects of mutations on the allosteric networks are ligand-specific, further underscoring that the dynamics of each complex are different and can be sorted by active vs inactive statuses.

Given that communication paths are simply chains of edges connecting two surfaces, we sought to determine whether a subset of edges would display weights correlating with EC_50_ of the receptor-ligand complexes. We compared the weights of all edges involved in ligand-H12 communication and a few edges emerged that satisfy this criteria (Figure 10), including the ASN35-ligand edge (ASN35_ligand). This edge was already identified in our analysis to be most prominent in the paths for all complexes, whether active or inactive. Future work will be needed to determine whether this correlation can be used as a predictive marker for ligand activation status.

**Figure 10:**
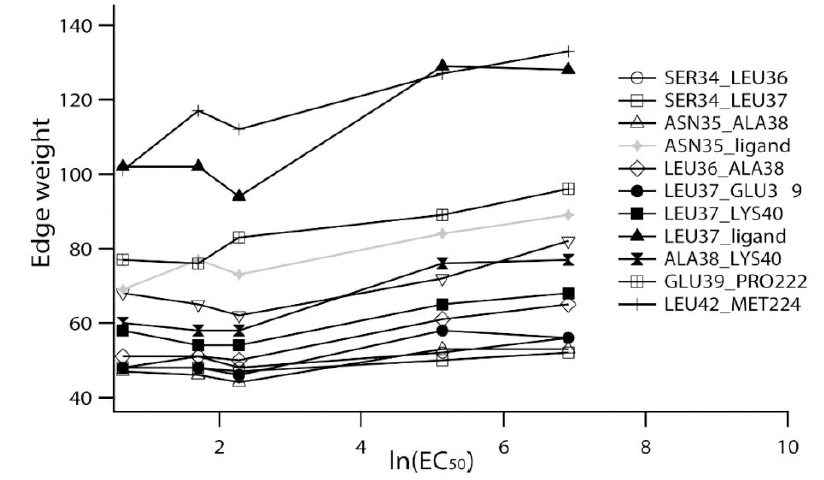
Edges show correlation with EC_50_. EC_50_s on x-axis in ascending order are for AncSR2-aldosterone (1.9 nM), AncSR2-progesterone (5.5 nM), AncSR2-cortisol (9.8 nM), AncSR2-DHT (172 nM) and AncSR2-estradiol (unmeasurable, 1000 nM used). Only edges for which r^2 > 0.8 are shown.

### Asn35, a conserved and crucial SR residue, is important for allostery in AncSR2

Evolutionarily conserved residues are believed to be integral to function and/or structure of a protein family. In addition to ASN35 being conserved across all naSRs (PR: ASN719, AR: ASN705, MR: ASN770, GR: ASN564), this residue has been demonstrated to be crucial for ligand-driven SR-activation in these SRs. This H3 asparagine residue contributes to PR activation by C17-hydroxylated progestins (65), anchors both androgens and nonsteroidal antiandrogens in the AR pocket (66), participates in hydrogen-bond networks that stabilize both ligands and H12 in MR (67) and interacts with C11/C21 OH groups of glucocorticoids (68). In all cases, mutation of ASN35 yielded deleterious effects on activation. The identification of this crucial residue and its role in AncSR2 allosteric communication is a striking result that validates the use of this computational method in identifying residues that both mediate allosteric communication and modulate protein function. Additionally, this result is consistent with work by Lockless at al which demonstrated in multiple protein families that evolutionarily conserved amino acids can link distant, coupled functional sites (28).

Interestingly, because ERs branched off directly from AncSR1, a threonine residue is found at this position in place of ASN in modern ERs (ERα: T347, ERβ: T299). However introduction of T347N in ERα did not have an effect on ligand response (69). Additionally, from work by Harms et al (61) it is known that this residue position does not distinguish A-ring aromatization in steroids. Therefore, the reduced strength observed in the ASN35-estradiol or ASN35-NPT edge is not necessarily the result of a mismatch or clash. However, it could be the indirect result of overall pocket destabilization. Of the 23 residues that line the SR ligand binding pocket, 12 differ between AncSR2 and ERα (Figure 11. shaded). Of the 12, up to 7 have been demonstrated to directly affect ligand binding in ERα upon mutagenesis (69-72). It is possible then that overall unfavorable conditions in the AncSR2 pocket for estrogen binding might be responsible for the reduced strength of the ASN35-ligand edge. More work will be needed to clarify this effect.

**Figure 11.**
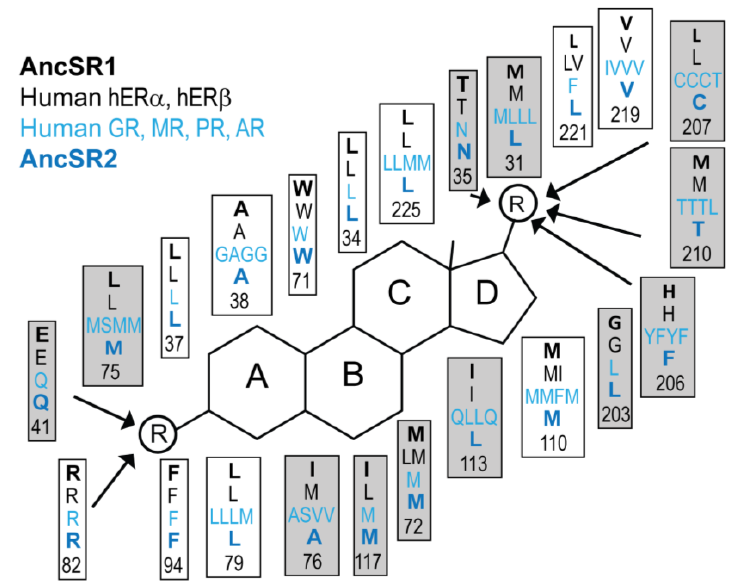
Binding pocket residues compared between ancient and extant SRs. (reproduced from (42)).

### Evolution of SR allosteric networks

SRs operate by a common mechanism, so presumably there is a commonality to the allosteric networks that facilitate activation in all SRs. Eick et al (42)suggested that promiscuity in AncSR2’s ligand responses resulted, in part, from the excess volume in the pocket cavity, as well as the presence of polar side chains within the pocket. These two features permit AncSR2 to respond to a range of ligands, some of which evolved after AncSR2 was already present (73). Combined with these ideas, our work suggests that a certain configuration of AncSR2 is induced in combination with activating ligands, but lost in inactive ligands. This configuration appears to be reflected in the allosteric networks studied here. These changes are subtle and do not reflect as large helical movements, but we are still able to capture these differences through the use of the dynamic network approach using correlation. (I.e. estrogen binding to AncSR2 does not place it in an optimal configuration for strong allosteric communication between the ligand and H12). These results underscore the utility of MD simulations in incorporating dynamics which permits subtle conformational changes to be sampled.

To fully elucidate the conformational and network effects observed here, allosteric networks of other SRs will need to be investigated with both activating and inactive ligands. One would expect to see that activating ligands achieve conformations, and thus networks, that are not observed with inactive ligands. Presumably, an inverse agonist would also need to induce communication between the LBP and H12, but in a manner that is distinct from agonists. But to understand the basis of selectivity in activation of SRs, further work will be needed to determine whether activating ligands themselves perturb networks uniquely in a promiscuous receptor. In other words, do progesterone and cortisol, both activating ligands, perturb AncSR2 networks in distinct manners? These studies are outside the scope of this particular work.

Ultimately, our goal is to determine whether the alterations in allosteric networks obtained from a computer simulation can be used to predict *a priori* the activation status of a ligand. Having determined that restorative mutations will alter *in silico* allosteric networks in an inactive ligand complex, strengthening LBP-AF-2 communication, it would be imperative to determine whether we can use mutational analysis to predict mutations that will either ablate ligand activity or introduce activity to a previously inactive ligand. Finally, allosteric networks in this study appear to correlate with ligand activation, quantified by EC_50_s. It is not clear at this time but will be interesting to determine whether information about other aspects of receptor-ligand interactions, including ligand-binding affinity, ligand efficacy, and receptor stabilization can potentially be extracted from these networks.

## Acknowledgments

This research was supported by NIH/NIGMS K12GM00680-18 (CDO) and a W.M. Keck Foundation Grant (EAO). We would like to thank Dr. Jen Colucci for editing contributions.

## Author Contributions

CDO and EAO conceived and designed the experiments. CDO performed experiments and analyzed the data. CDO and EAO wrote the paper.

## Supporting Information

**Table S1.** Edges in top 2000 paths for each AncSR2-hormone complex. Edges which appeared in at least 15% (of any AncSR2-ligand complex) are shown; values in boxes denote the percentage of the edge for each complex. Shaded boxes indicate edges which appear in at least 25%, while highlighted (yellow) indicates the ASN35-ligand edge which is the most prevalent edge across all simulations.

|  | DHT | ALD | COR | PROG | EST |
| --- | --- | --- | --- | --- | --- |
| ILE228, LEU225 | 25.1 | 26.35 | 22.55 | 17.85 | 27.3 |
| LEU225, ASN35 | 19.9 | 10.35 | 12.95 | 17.7 | 10.75 |
| ASN35, Ligand | 33.7 | 40.05 | 34.6 | 45.2 | 31.5 |
| ALA38, Ligand | 25.35 | 14.4 | 25.05 | 29.9 | 19.95 |
| ILE228, VAL226 | 28.2 | 25.45 | 26.45 | 22.55 | 24.45 |
| VAL226, PRO222 | 13.8 | 16.35 | 17.2 | 15.6 | 18 |
| PRO222, ASN35 | 17.35 | 13.5 | 17.5 | 22.25 | 17.15 |
| LEU225, PRO222 | 14.1 | 14.05 | 17.55 | 16 | 16.8 |
| PHE221, ASN35 | 15.3 | 13.9 | 16.7 | 19.45 | 14.8 |
| ILE228, MET224 | 14.2 | 12.6 | 12.6 | 9.5 | 15.5 |
| ASN35, ALA38 | 14.45 | 0 | 15.9 | 16.3 | 14.25 |
| LEU34, Ligand | 31.95 | 20.95 | 17.4 | 12.8 | 26.2 |
| LEU225, GLU227 | 24 | 22.9 | 23.2 | 27.55 | 21.35 |
| ILE228, ALA231 | 12 | 11.3 | 14.6 | 21.05 | 11.5 |
| VAL226, ILE229 | 17 | 19.5 | 0 | 19.85 | 0 |
| ALA231, ILE229 | 8.3 | 8.45 | 11.05 | 17.6 | 8.3 |
| GLN232, ILE229 | 8 | 10.7 | 11.9 | 15.65 | 8.45 |
| ILE229, GLU227 | 11.1 | 12.15 | 10.3 | 16.15 | 8.15 |

**Figure S1.**
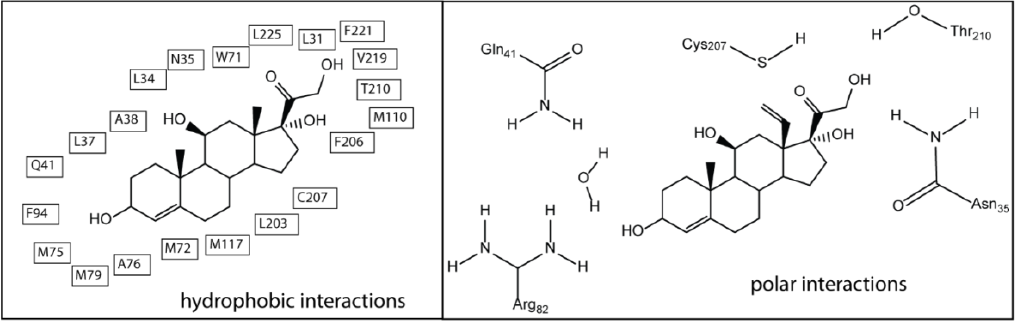
Binding pocket interactions in AncSR2. Hydrophobic interactions (defined by C-C atom distances) and polar interactions as observed in simulations are shown, using cortisol.

**Table S2.**
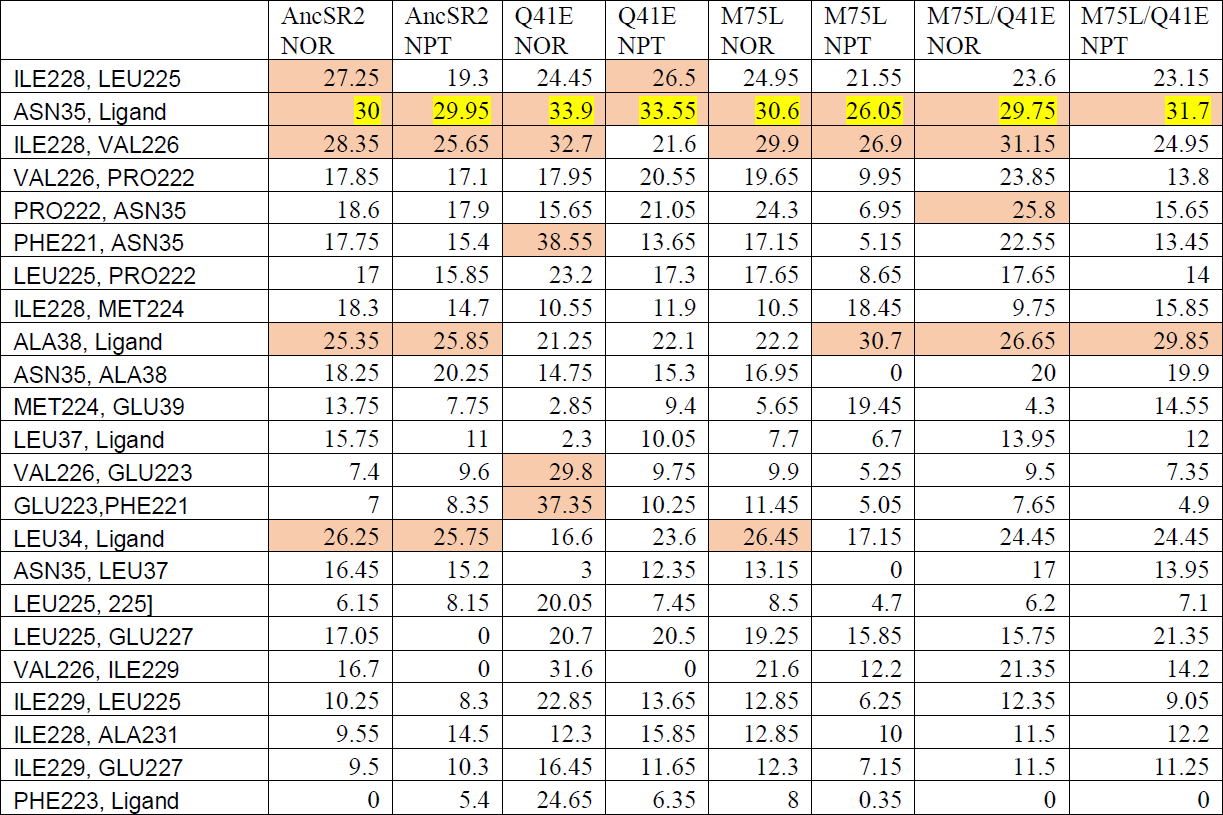
Edges in top 2000 paths for each AncSR2-hormone complex. Edges which appeared in at least 15% (of any AncSR2-ligand complex) are shown; values in boxes denote the percentage of the edge for each complex. Shaded boxes indicate edges which appear in at least 25%, while highlighted (yellow) indicates the ASN35-ligand edge which is the most prevalent edge across all simulations.

|  | AncSR2<br>NOR | AncSR2<br>NPT | Q41E<br>NOR | Q41E<br>NPT | M75L<br>NOR | M75L<br>NPT | M75L/Q41E<br>NOR | M75L/Q41E<br>NPT |
| --- | --- | --- | --- | --- | --- | --- | --- | --- |
| ILE228, LEU225 | 27.25 | 19.3 | 24.45 | 26.5 | 24.95 | 21.55 | 23.6 | 23.15 |
| ASN35, Ligand | 30 | 29.95 | 33.9 | 33.55 | 30.6 | 26.05 | 29.75 | 31.7 |
| ILE228, VAL226 | 28.35 | 25.65 | 32.7 | 21.6 | 29.9 | 26.9 | 31.15 | 24.95 |
| VAL226, PRO222 | 17.85 | 17.1 | 17.95 | 20.55 | 19.65 | 9.95 | 23.85 | 13.8 |
| PRO222, ASN35 | 18.6 | 17.9 | 15.65 | 21.05 | 24.3 | 6.95 | 25.8 | 15.65 |
| PHE221, ASN35 | 17.75 | 15.4 | 38.55 | 13.65 | 17.15 | 5.15 | 22.55 | 13.45 |
| LEU225, PRO222 | 17 | 15.85 | 23.2 | 17.3 | 17.65 | 8.65 | 17.65 | 14 |
| ILE228, MET224 | 18.3 | 14.7 | 10.55 | 11.9 | 10.5 | 18.45 | 9.75 | 15.85 |
| ALA38, Ligand | 25.35 | 25.85 | 21.25 | 22.1 | 22.2 | 30.7 | 26.65 | 29.85 |
| ASN35, ALA38 | 18.25 | 20.25 | 14.75 | 15.3 | 16.95 | 0 | 20 | 19.9 |
| MET224, GLU39 | 13.75 | 7.75 | 2.85 | 9.4 | 5.65 | 19.45 | 4.3 | 14.55 |
| LEU37, Ligand | 15.75 | 11 | 2.3 | 10.05 | 7.7 | 6.7 | 13.95 | 12 |
| VAL226, GLU223 | 7.4 | 9.6 | 29.8 | 9.75 | 9.9 | 5.25 | 9.5 | 7.35 |
| GLU223,PHE221 | 7 | 8.35 | 37.35 | 10.25 | 11.45 | 5.05 | 7.65 | 4.9 |
| LEU34, Ligand | 26.25 | 25.75 | 16.6 | 23.6 | 26.45 | 17.15 | 24.45 | 24.45 |
| ASN35, LEU37 | 16.45 | 15.2 | 3 | 12.35 | 13.15 | 0 | 17 | 13.95 |
| LEU225, 225] | 6.15 | 8.15 | 20.05 | 7.45 | 8.5 | 4.7 | 6.2 | 7.1 |
| LEU225, GLU227 | 17.05 | 0 | 20.7 | 20.5 | 19.25 | 15.85 | 15.75 | 21.35 |
| VAL226, ILE229 | 16.7 | 0 | 31.6 | 0 | 21.6 | 12.2 | 21.35 | 14.2 |
| ILE229, LEU225 | 10.25 | 8.3 | 22.85 | 13.65 | 12.85 | 6.25 | 12.35 | 9.05 |
| ILE228, ALA231 | 9.55 | 14.5 | 12.3 | 15.85 | 12.85 | 10 | 11.5 | 12.2 |
| ILE229, GLU227 | 9.5 | 10.3 | 16.45 | 11.65 | 12.3 | 7.15 | 11.5 | 11.25 |
| PHE223, Ligand | 0 | 5.4 | 24.65 | 6.35 | 8 | 0.35 | 0 | 0 |

